# Grid cells encode reward distance during path integration in cue-rich environments

**DOI:** 10.1101/2025.09.03.674124

**Authors:** Satoshi Kuroki, Sébastien Royer

## Abstract

The medial entorhinal cortex supports both path integration and landmark anchoring, but how these computations interact during goal-directed navigation is unclear. We show that grid cells dissociate from landmarks and instead encode reward distance when mice perform a path integration task on a cue-rich treadmill. Grid cell population activity reset at rewards and shifted coherently across trials, consistent with continuous attractor dynamics realigned by rewards. Furthermore, grid cells exhibited reduced spatial scales, broadened theta frequency distributions, and altered temporal coordination. These phenomena were captured by a theta interference model incorporating cell competition and two sets of theta oscillating inputs whose frequencies shifted apart. Switching to cue-based navigation stabilized the firing fields and partially restored grid scale, theta frequencies and temporal structure. These results demonstrate that MEC circuits flexibly reset to encode goal-directed trajectories, and suggest that continuous attractor and interference mechanisms normally cooperate but can decouple under path integration demands.

## Introduction

The medial entorhinal cortex (MEC) is a key structure for spatial navigation, supporting both path integration and landmark anchoring^1-5^. Yet the mechanisms by which these computations emerge and interact to guide goal-directed navigation remain unclear.

Grid cells, with their strikingly periodic firing fields organized into modules of distinct scale and orientation, have been proposed to implement path integration^1,6-9^, but how their regularity arise has been debated. Competing models attribute grid periodicity to continuous attractor dynamics^10^ or to interference between theta oscillatory inputs^11^. The attractor framework emphasizes recurrent network interactions that preserve grid relationships across environments and states^12-15^, whereas the interference model predicts that changes in theta input frequency alter grid scale and phase precession. A coexistence of these mechanisms has been proposed^16^, but direct experimental evidence remains limited.

A second unresolved question is whether grid cells directly support goal-directed navigation, for example by encoding goal distance^17^, or whether they primarily implement landmark-referenced path integration. Although grid fields are typically anchored to environmental landmarks^1^, they can be influenced by reward locations^18^ and rapidly re-anchored through plasticity mechanisms^19-20^, suggesting a flexible alignment. Disentangling whether grid activity reflects landmarks, rewards, or internally computed distances has been challenging, since rewards usually occur at fixed positions.

To address this, we developed a path integration task in which mice received rewards after fixed running distances but at varying environmental locations. Using silicon probe recordings in the MEC, we compared this task to open-field foraging and a cued navigation condition with fixed reward locations. We found that grid and border cells disengaged from landmarks and instead repeated activity patterns aligned to reward distance. Grid fields also exhibited reduced scale and broader theta frequency distributions, consistent with a theta interference model in which the two input frequencies shifted apart. These findings demonstrate that MEC circuits can flexibly reset to represent goat-directed trajectories, and suggest that continuous attractor and interference mechanisms normally cooperate to stabilize spatial maps but decouple under the atypical requirement of path-integration in a cue-rich environment.

## Results

### Behavior

Mice (n = 4) were trained to forage in an open field and to perform two tasks on a cue-rich treadmill: a path integration (PI) task, in which water rewards were delivered after a fixed 105-cm running distance at variable belt positions, and a cue task, in which water rewards were delivered at two fixed, unevenly spaced locations (Fig.1a). Following 17-31 training days, mice displayed robust anticipation of rewards, evidenced by increased lick rated and reduced running speed before reward delivery (Fig. 1b).

**Figure 1.**
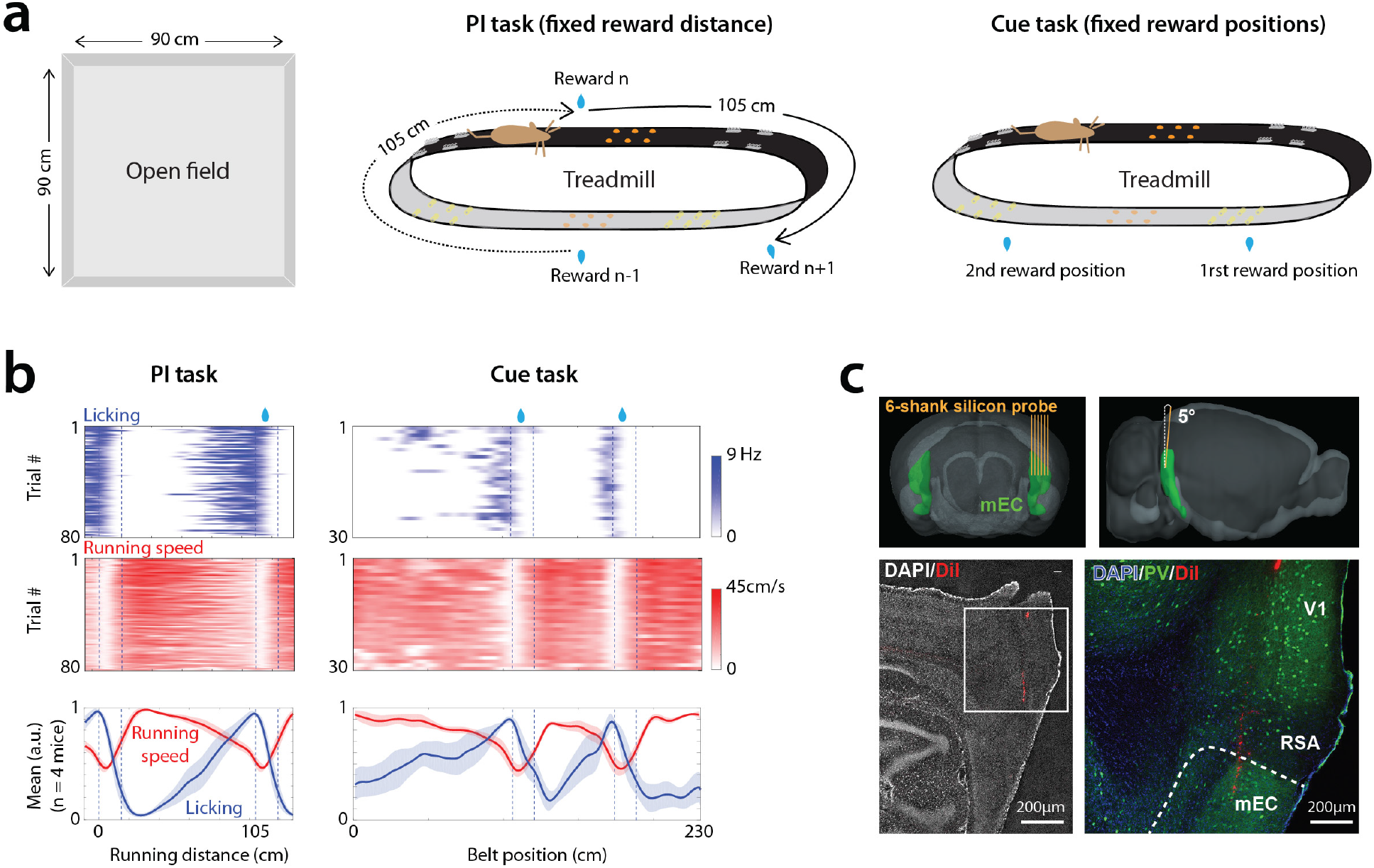
Behavior paradigm and silicon probe recording in medial entorhinal cortex (MEC). a, Schematic of the open-field arena (left) and treadmill apparatus with reward delivery protocols for the path integration (PI) and cue tasks (right). b, Example session showing licking (blue) and running speed (red) across individual trials (color-coded) and session averages (traces), aligned to reward distance (PI task, left) or belt position (cue task, right). c, Top, 3D renderings of a mouse brain (Allen Brain Atlas) showing the target MEC and probe insertion angle (coronal, left; lateral). Bottom, histology from an example mouse showing the probe track (DiI, red) within MEC (DAPI, blue). Right, higher magnification with PV staining (green) marking MEC layer 2/3.

### Identification of grid and border cells

We chronically implanted silicon probes in MEC (Fig. 1c) and recorded in both open field and treadmill on each day. Across 4-6 days per mouse, we isolated 1421 cells and tracked them across 2-3 consecutive sessions per day (11 open field/PI, 4 open field/cue and 4 open field/PI/cue). Grid cells (n = 216) and border cells (n = 25) were identified in the open field using established grid and border score criteria^2,21^. To account for the prospective coding of grid cells^22^, rate maps were computed with optimal prospective shifts. We then analyzed grid and border cell activity on the treadmill by constructing firing rate maps for individual belt cycles and for reward-to-reward journeys (105 cm), referred to here as trials.

### Grid and border cells dissociate from landmarks during path integration

During the PI task, grid cells displayed multiple fields that failed to align across belt positions in contrast to their consistent alignment with landmarks during the cue task (Fig. 2a-b, Supplementary Fig. 1). Correlation of firing rate maps across trials were significantly higher in the cue task than in the PI task (Fig. 2c). In the PI task, however, many grid cells aligned consistently with reward distance, and correlations across reward-to-reward journeys were significantly higher than across belt cycles, indicating stronger encoding of reward distance than belt position (Fig. 2a-c). Border cells showed a similar dissociation from landmarks in the PI task (Supplementary Fig. 2), with higher stability in the cue task (Fig 2c). Thus, both grid and border cells anchored reliably to landmarks during the cue task but disengaged from landmarks in the PI task, with grid cells preferentially encoding reward distance.

**Figure 2.**
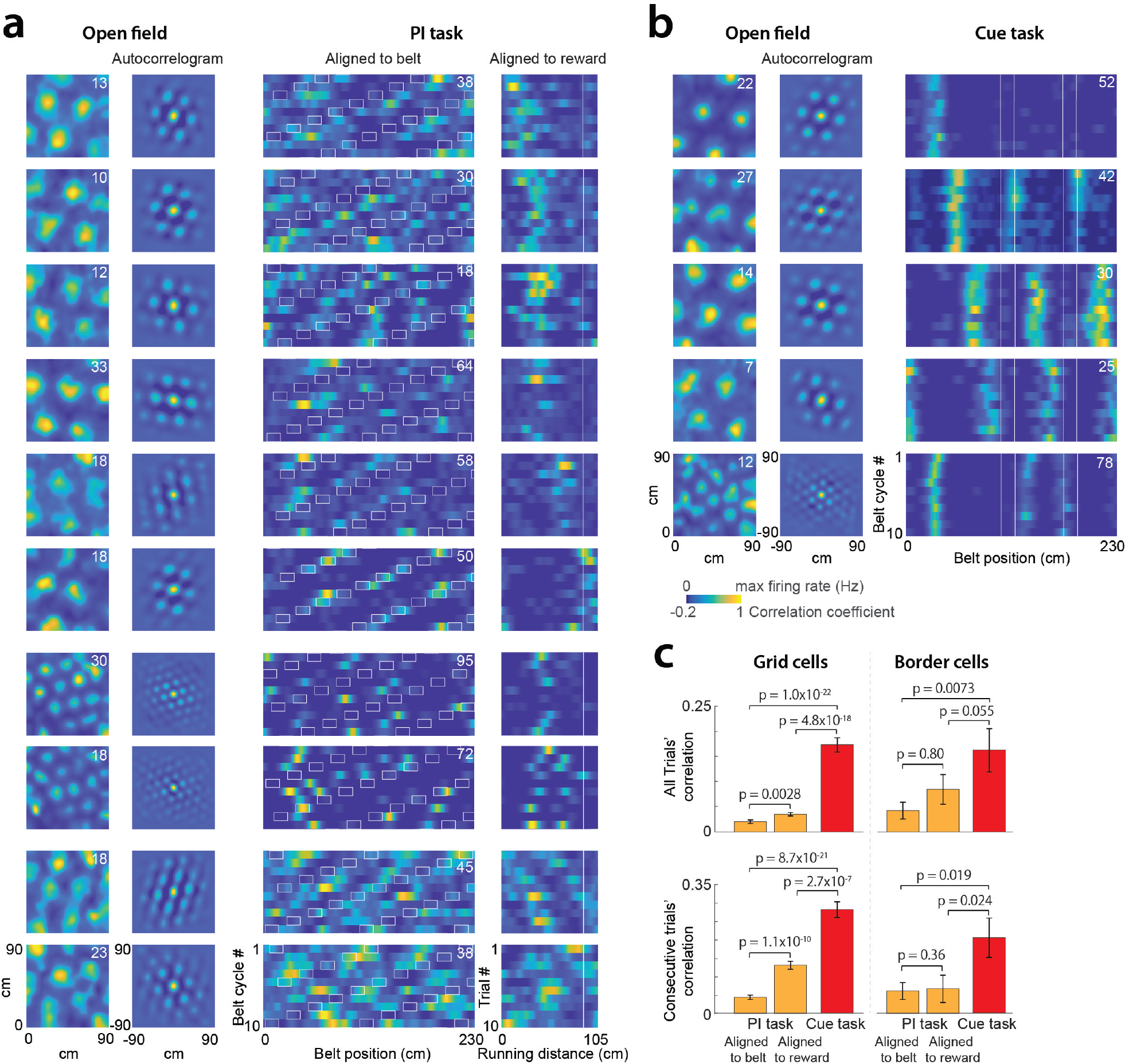
Grid cell firing in open field, path integration (PI) and cue tasks. a, Activity of example grid cells recorded consecutively in open field and PI task (one cell is represented in each row). First and second columns: rate maps and spatial autocorrelograms in open field. Third and fourth columns: rate maps during the PI task, organized by belt cycles (third) or journeys to reward (fourth). White rectangles indicate reward locations. Top six examples show cells with comparable grid sizes encoding different reward distances; next two show cells with shorter grid sizes; bottom two show cells without position or reward distance coding. b, Same as in a for grid cells recorded consecutively in open field and cue task. c, Trial-by-trial correlations of rate maps (mean±s.e.m., grid cells: n = 155 PI, 102 Cue; border cells: n = 17 PI, 15 Cue). Comparisons include all trial pairs (top) and consecutive trial pairs (bottom), for PI (orange) and cue (red) tasks (Wilcoxon signed-rank and rank-sum tests).

### Grid cell population vectors reset at rewards

To assess how rewards influenced population activity, we computed grid cell population vectors (PVs) for each trial and quantified similarity across trials using inter-trial correlation matrices (Fig. 3a). Hierarchical clustering revealed clusters of trials with highly similar PVs (Fig. 3b). PV similarity was not determined by trial start position (Fig. 3c) or session period (Fig. 3d), but correlated significantly with behavioral similarity (lick rate and running speed; Fig. 3g). Similar PVs often recurred in successive trials more than expected by chance (Fig. 3e-f). These findings indicate that reward delivery resets grid cell PVs, with variability paralleling behavioral variability.

**Figure 3.**
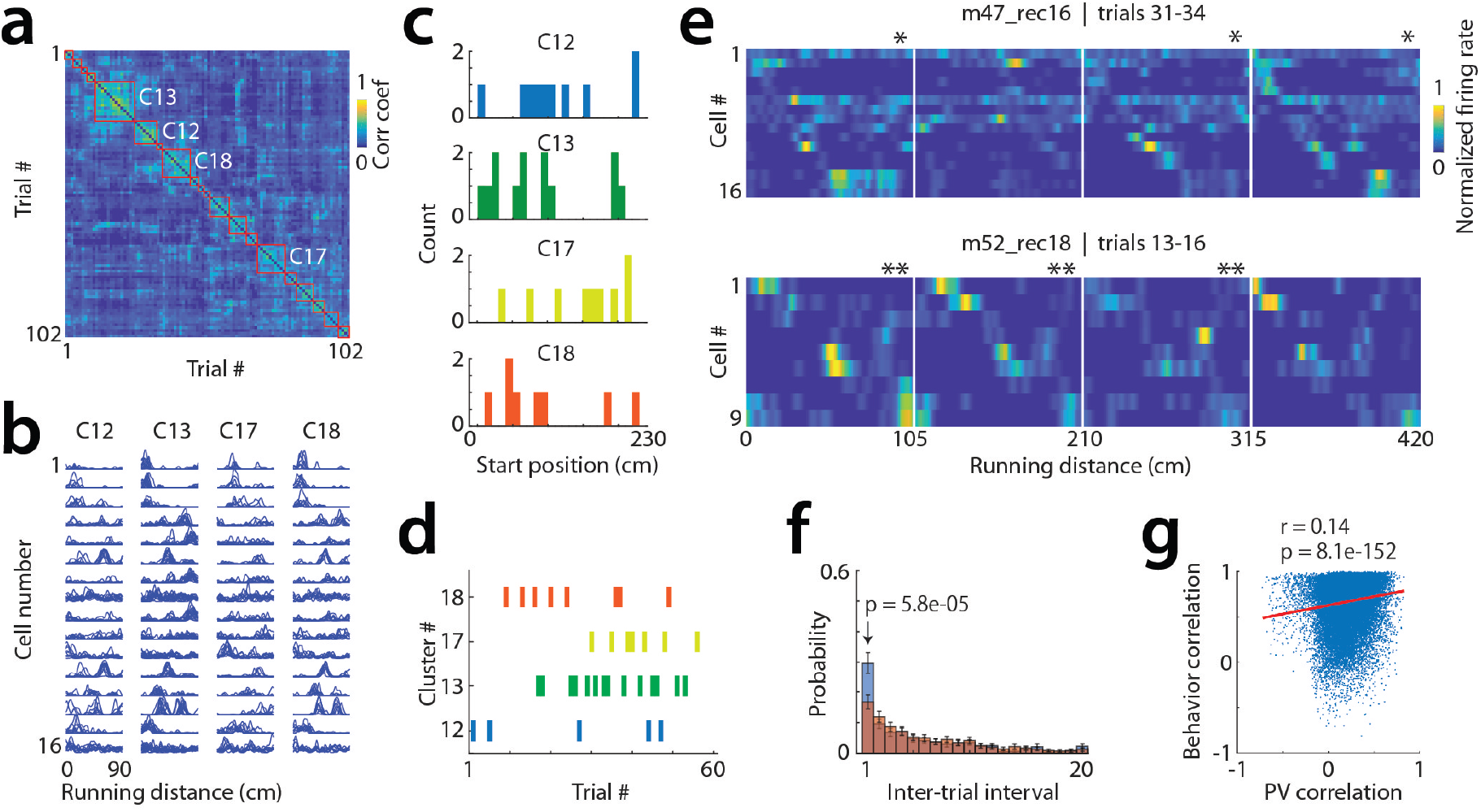
Spatiotemporal organization of grid cell activity during the PI task. a, Correlation matrix of population vectors across trials, sorted by similarity. Vectors include firing rates of all simultaneously recorded grid cells across spatial bins of reward-directed journeys. b, Overlay of firing rates from individual trials within the four clusters highlighted in a. c, Start positions of trials belonging to each cluster, distributed across the full belt. d, Example grid cell activity across consecutive trials in two mice. Stars mark trials with similar population vectors. e, Trial occurrence times (trial numbers) for the four clusters in a. Clusters are distributed across the session and sometimes occur in succession. f, Distribution of inter-trial intervals for real clusters (blue, 15 sessions, 4 mice) and shuffled clusters (orange, one shuffle per session; n = 65 clusters; Wilcoxon signed-rank test). g, Correlation between behavioral vectors (running speed and licking) and grid cell population vectors (n = 36,993 trial pairs; Pearson correlation).

### Grid cells shift coherently across trial clusters

Comparisons across trial clusters revealed coherent shifts in grid cell activity (Fig. 4a). Spatial cross-correlograms of PVs indicated a broad distribution of phase lags across clusters (Fig. 4b–c), while pairwise cross-correlograms demonstrated that relative spatial relationships between grid cells were preserved (Fig. 4d–f), including across PI and cue tasks (Supplementary Fig. 3). Simulations of linear trajectories through 2D grid maps^23^ showed that only collinear shifts reproduced both the broad distribution of PV shifts and the preservation of cell–cell spatial relationships observed experimentally (Supplementary Fig. 4). Thus, grid cells undergo coherent collinear shifts across trial clusters.

**Figure 4.**
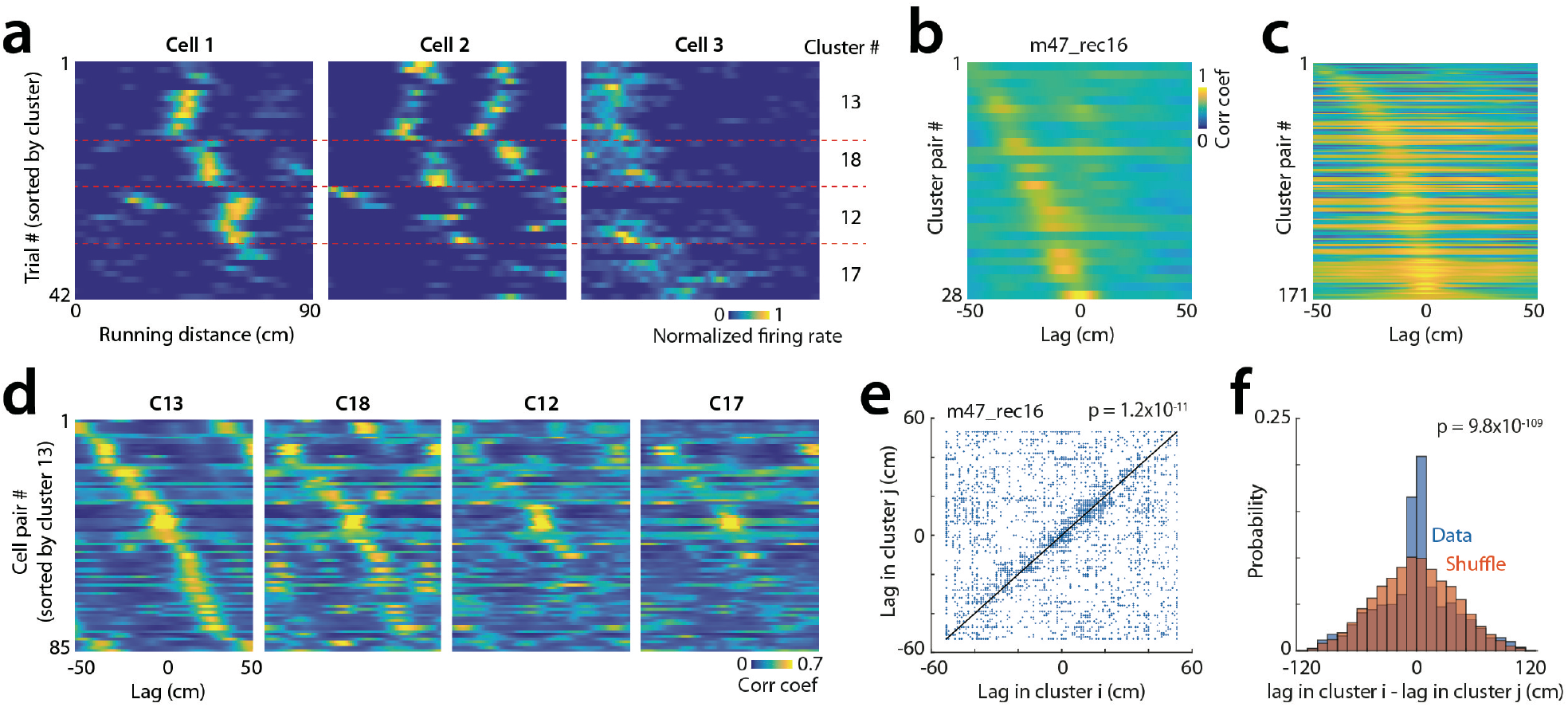
Coherent spatial shifts of grid cells. a, Firing rate maps of three grid cells, organized by clusters (as in Fig. 3a). Note spatial shifts across clusters. b, Spatial cross-correlograms of population vectors between cluster pairs, sorted by peak position, for one session. c, Same as b, pooled across all sessions and mice. d, Spatial cross-correlograms between cell pairs within clusters (four clusters from Fig. 3a). Pairs are ordered by peak position in cluster C13 and preserved across clusters, showing stable cell–cell relationships. e, Cross-cluster comparison of spatial shifts between cell pairs (n = 14,209 cluster pairs; Pearson correlation). f, Changes in spatial shifts across clusters (blue) versus shuffled cell identities (orange, one shuffle per cluster; n = 14,209 pairs; Wilcoxon rank-sum test on absolute values).

### Grid size is reduced and coherence weakened in the PI task

Grid size on the treadmill is expected to be similar or larger than in the open field, according to CAN predictions for linear environments^23^. Unexpectedly, grid sizes estimated from inter-field distances were smaller during the PI task compared to the open field (Fig. 5a–e), whereas grid size in the cue task did not differ (Fig. 5e). Reduced grid size was accompanied by weaker coherence among grid cells (Fig. 5f–g), particularly for cells with smaller grids. These findings suggest that grid periodicity during the PI task cannot be explained solely by CAN dynamics.

**Figure 5.**
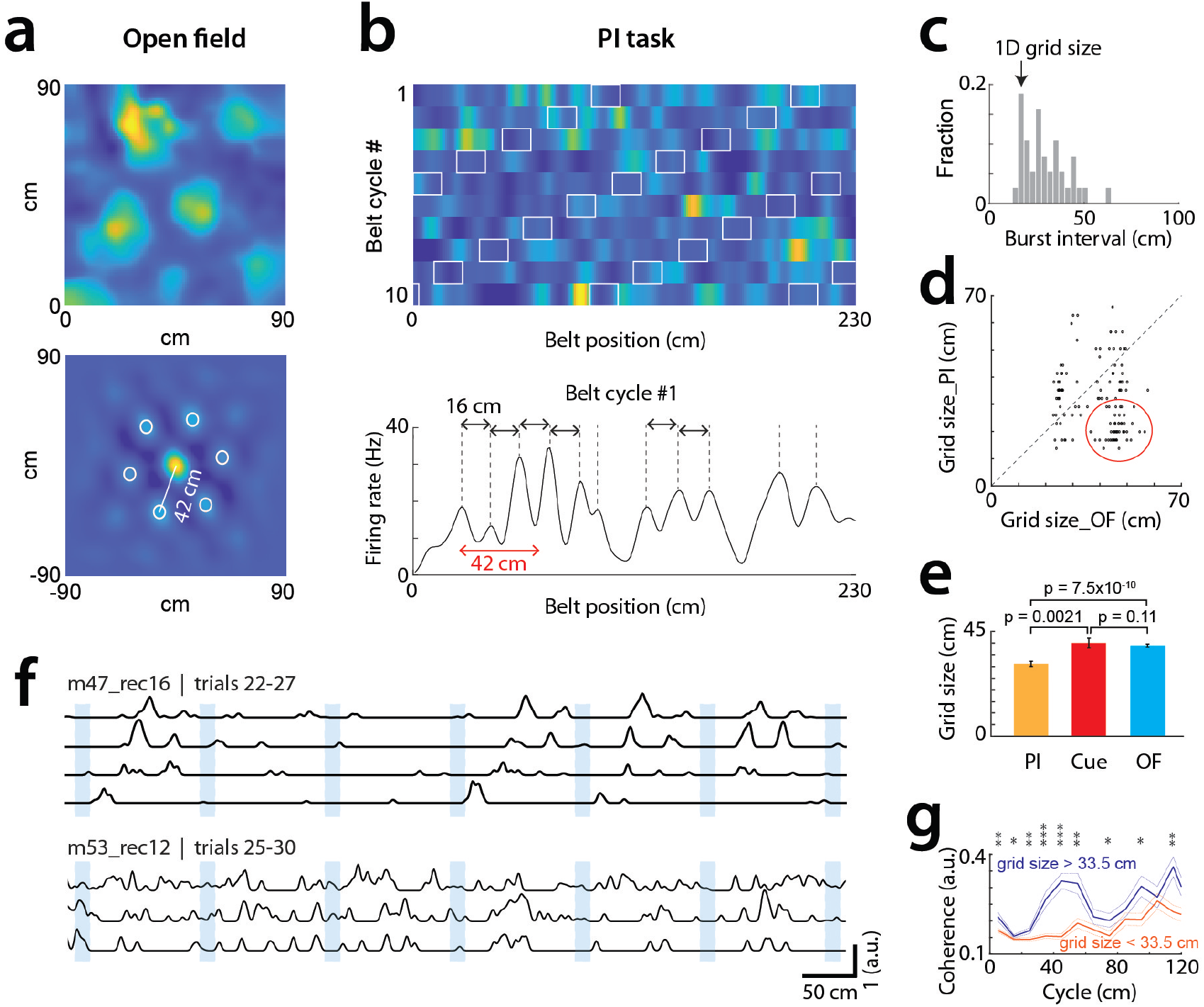
Grid size reduction during the PI task. a, Example grid cell in open field: rate map (top) and autocorrelogram (bottom), grid size = 42 cm. b, Same cell during PI task: firing rates across multiple belt cycles (top) and one belt cycle (bottom). Black and red arrows mark 16 cm (1D burst interval) and 42 cm (2D grid size), respectively. c, Distribution of inter-burst intervals for the cell in b; 1D grid size defined as peak. d, Grid size in PI versus open field. Red circle highlights cells with substantially smaller grid size in PI. e, Grid size across conditions (mean ± s.e.m., n = 134 PI, 100 Cue, 189 OF cells; Wilcoxon rank-sum test). f, Two session examples with co-recorded grid cells of large (top) and small (bottom) grid sizes. Large grid cells show coherent activation patterns across trials, unlike small grid cells. g, Coherence across grid cell pairs in PI, separated by grid size (large: >33.5 cm, blue, n = 45 pairs; small: <33.5 cm, red, n = 78 pairs; Wilcoxon rank-sum test). *p < 0.05, **p < 0.01, ***p < 0.001.

### Grid cells exhibit broader theta frequency ranges in the PI task

Because oscillatory interference models predict a relationship between theta frequency differences and grid size^11^, we examined theta rhythmicity. Mean firing rates did not differ between treadmill tasks, although they were higher in the open field (Supplementary Fig. 5). Both LFPs and grid cells exhibited theta rhythmicity (Fig. 6a–f), with strongest rhythmicity in the open field. Theta frequencies were lowest in the cue task and highest in the open field, but most notably, the spread of theta frequencies across grid cells increased markedly in the PI task (Fig. 6f; Supplementary Fig. 6).

**Figure 6.**
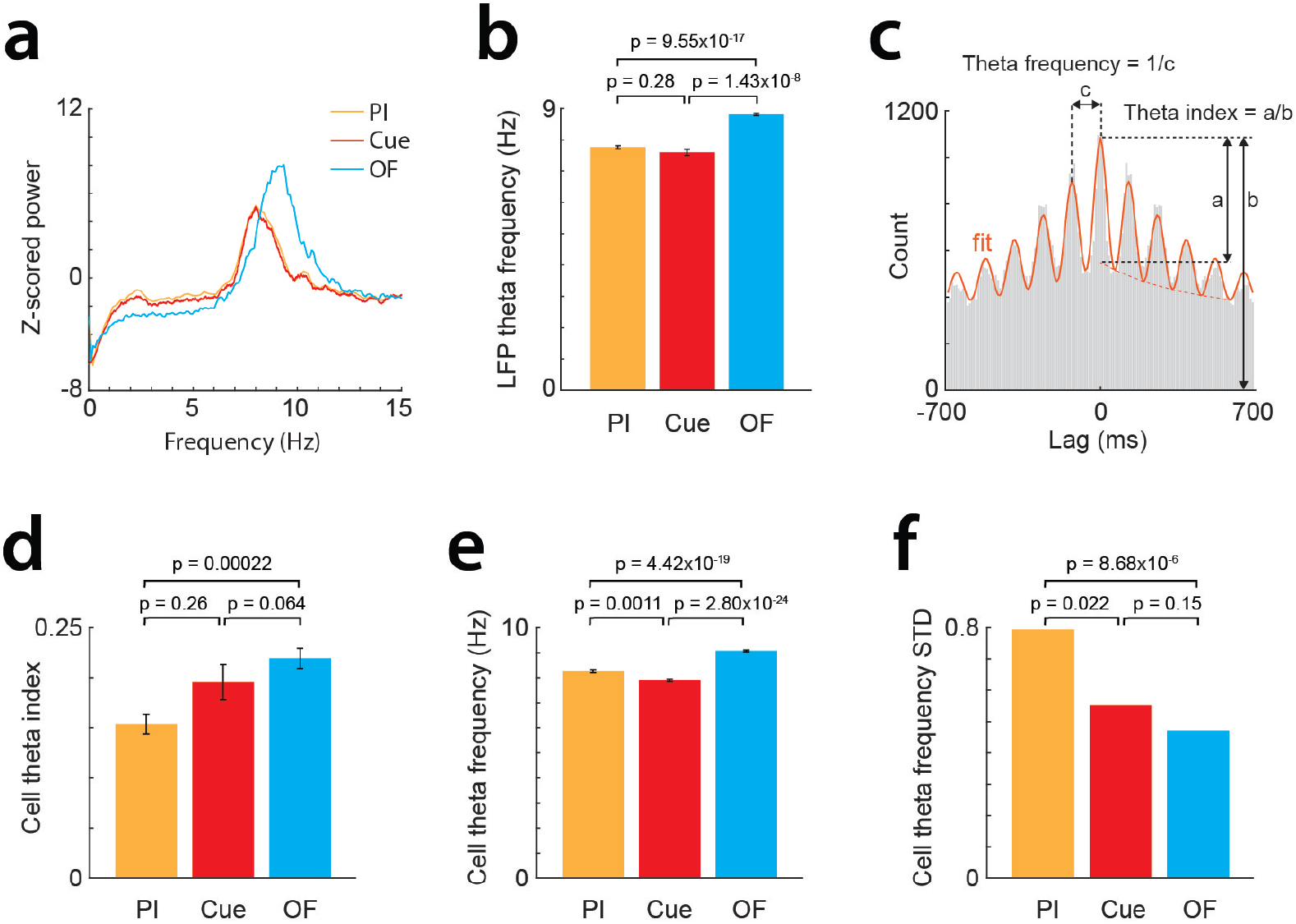
Altered theta oscillations during the PI task. a, Example LFP power spectra recorded consecutively in open field, PI, and cue tasks. b, Peak theta frequency across conditions (mean ± s.e.m., n = 15 PI, 8 Cue, 19 OF sessions; Wilcoxon rank-sum test). c, Spike-time autocorrelogram of a grid cell (gray), sinusoidal fit (red), and definitions of theta index and frequency. d, Theta index across conditions (mean ± s.e.m., n = 96 PI, 65 Cue, 151 OF cells; Wilcoxon rank-sum test). e, Theta frequency across conditions (same cells as in d). f, Standard deviation of theta frequency across conditions (same cells; Levene’s test).

### Theta interference model accounts for frequency and grid size changes

To test whether altered theta frequencies could explain reduced grid size, we simulated a simple interference model in which grid cells received excitatory input from two groups of oscillating cells at distinct theta frequencies and competed via feedback inhibition (Fig. 7a). The model produced theta rhythmicity and beat patterns resembling grid periodicity. Increasing the gap between the two input frequencies simultaneously reproduced the shorter grid sizes and broader theta frequency distributions observed in the PI task (Fig. 7b–f). This supports a role for theta interference in shaping grid cell periodicity.

**Figure 7.**
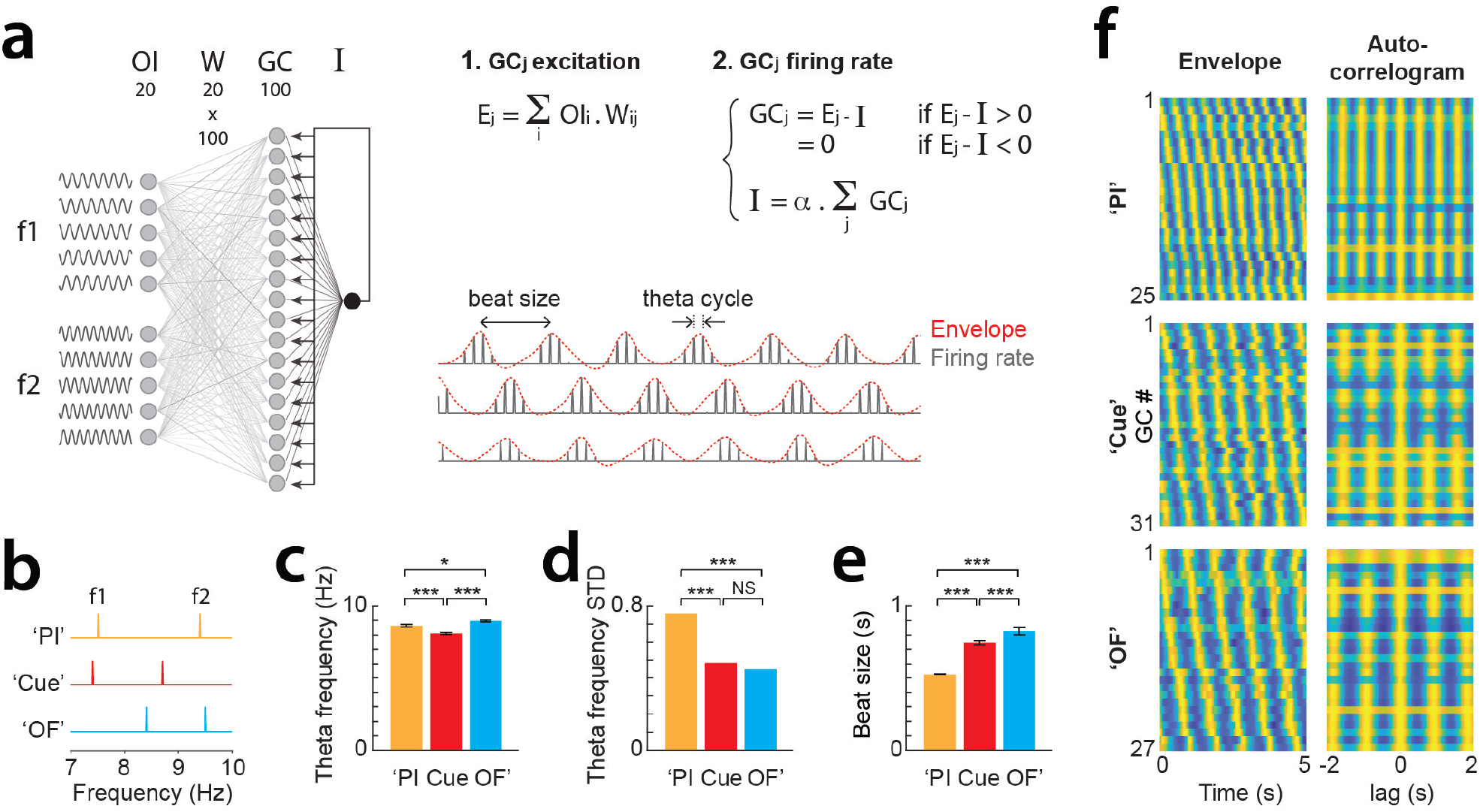
Interference model reproduces theta frequency and grid size alterations. a, Left, model schematic: 100 grid cells (GCs) receive random inputs from 20 oscillatory cells (OCs) with varying theta phases, divided into low- and high-frequency groups (f1, f2). GCs also receive feedback inhibition (I). Upper right, steps of one model iteration: (1) excitation to GCj; (2) GCj activation and inhibitory feedback, with I estimated numerically. Lower right, firing rate traces of three example GCs, showing beat size and theta cycle. b–e, OC frequency settings reproducing grid cell theta frequencies (c), variability (d), and grid size alterations (e). *p < 0.05, ***p < 0.0005 (Wilcoxon rank-sum for c, e; Levene’s test for d). f, GC envelopes (left) and autocorrelograms (right) for the three OC frequency settings.

### Altered short-timescale interactions during the PI task

Finally, we examined spike-time cross-correlograms (CCGs) of grid cell pairs. While overall CCG profiles were preserved across tasks (Fig. 8a–b), short-latency interactions varied, with central peaks or troughs differing across conditions. A short-timescale interaction index (STSI; Fig. 8c) revealed larger changes between PI and the other tasks than between cue and open field (Fig. 8d–e). These results suggest that while long-range grid cell interactions remain intact, short-timescale dynamics are selectively altered in the PI task.

**Figure 8.**
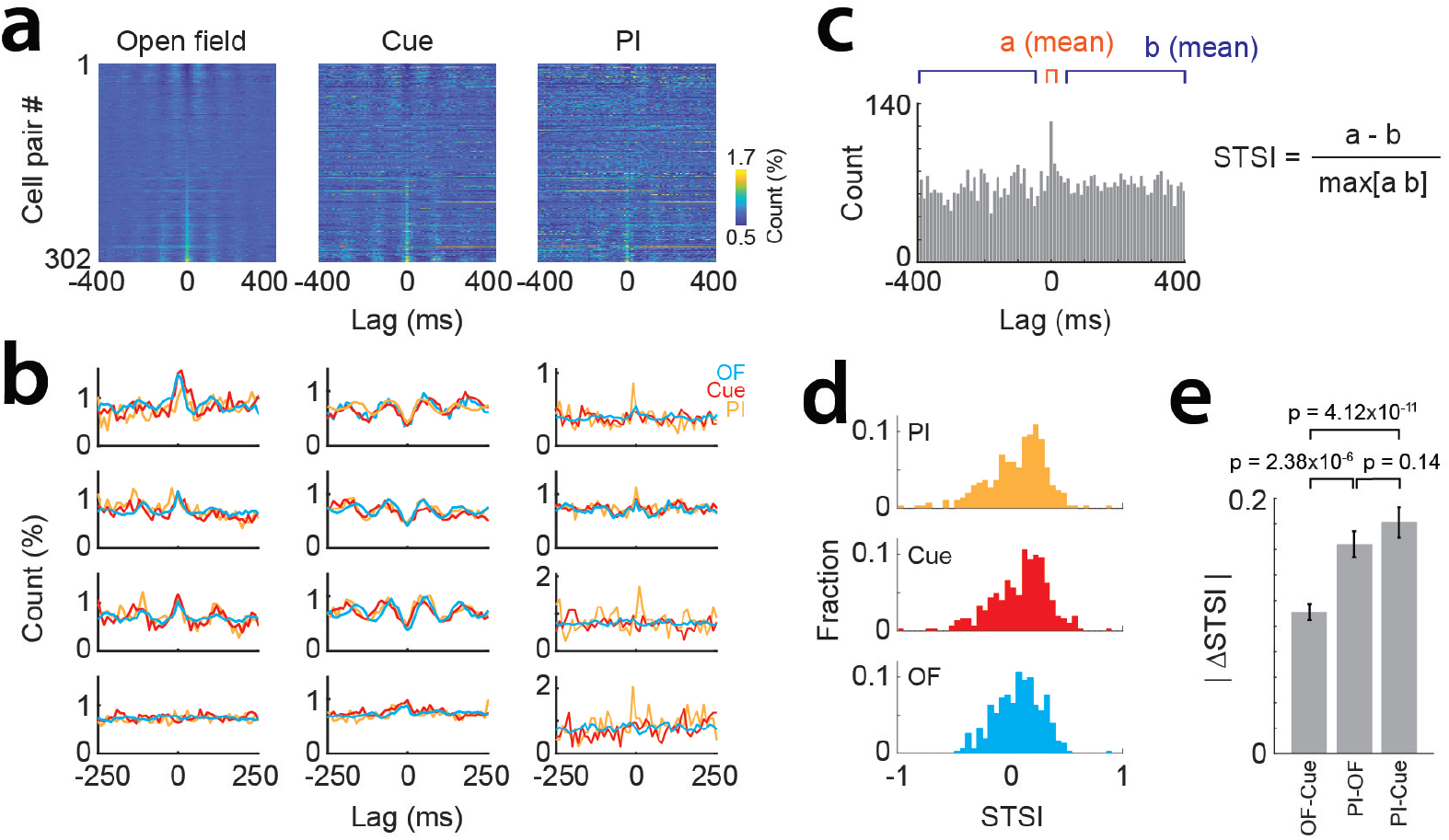
Short-timescale interactions between grid cells. a, Spike-time cross-correlograms (CCGs) between grid cell pairs recorded in open field, PI, and cue tasks. Pairs sorted by open field peak positions. Patterns from open field largely preserved in other tasks. b, Overlays of CCGs across tasks. Each of the first two columns: one reference cell with multiple partners. Note preserved task-specific CCG profiles but variability across partners. Rightmost column: examples of altered short-timescale interactions (STSI) during PI. c, Definition of STSI. d, STSI distributions across conditions. e, Changes in STSI across conditions (mean ± s.e.m. of absolute values, n = 302 pairs; Wilcoxon signed-rank test).

## Discussion

Grid cells have been studied across diverse environments and behavioral conditions^1,18,22,24-27^. Here we provide the first characterization of their activity patterns in a cue-rich treadmill, previously used to probe place cells in hippocampus and retroplenial cortex, and examine the special case of path integration-based navigation in a cue-rich environment.

Environmental landmarks and boundaries typically exert strong control over MEC activity, with border and object-vector cells mapping their proximity^2,4^ and grid cells stably anchoring to environmental features and reseting on the repeated alleys of a hair pin maze^24^ or at the ends of a linear track^28^. In circular tracks lacking landmarks, grid cells fail to maintain stable firing fields and instead encode travel distance^25^ as in treadmills^26^. In our cue task, the landmarks of the belt likely stabilized grid activity, providing inputs to correct accumulated errors and potentially reseting the CAN attractor on each belt cycle. Strikingly, however, during the path integration task both grid cells and border cells dissociate from landmarks. This finding aligns with recent reports of rapid grid realignment during self-motion-based navigation in open arenas^20^, supporting the idea that grid anchoring to landmarks is relatively weak and malleable. While landmarks did not reset grid cells in our task, they may still have contributed motion-related inputs to support path integration, as suggested in circular-track recordings^25^.

A longstanding hypothesis is that grid cells compute goal distance^17^. Our findings support this view: grid cells tended to reset at reward locations such that the same population code mapped multiple reward journeys. This provides a straightforward mechanism for estimating reward distance. For instance, grid ensembles at journey endpoints could recruit hippocampal reward cells^29^, while hippocampal reward cell activity could in turn reset the grid network to its starting state. The grid network might then evolve solely via CAN dynamics as mice progress to the next rewards. Such reward-based resetting may explain the directionality of grid codes in 1D tracks with rewards delivered at the ends of the track^28^ and the realignment of grid cells during goal direct navigation in open arenas^20^. In contrast, grid activity lacks such resetting during random foraging in open fields^1^ and circular tracks^25^ or during forced running on motorized treadmills^26^.

We also observed changes in grid cell theta oscillations across conditions. Previous work has shown lower theta frequency in novel compared to familiar environments^30^, in linear tracks versus open fields^25^ and in virtual reality relative to real world environments^31^. Our finding of a lower theta frequency on the treadmill than in the open field is consistent with both linear-track and virtual-reality effects. More strikingly, theta frequency varied much more across grid cells in the path integration task. Our modeling suggested that this effect resulted from an increased divergence between the two theta frequencies driving grid cells. High frequency (type 1) theta is associated with locomotion and path integration^32-34^ and accordingly modulated by running speed^32,34^. In this respect, the shift toward higher theta frequency from the cue to the path integration task could reflect the increased requirement of path integration. Similarly, the higher theta frequency in open field relative to 1D environments (ref25 and current results) and in real world relative to virtual reality^31^ might originate from a larger involvement of path integration mechanisms. By contrast, low frequency (type 2) theta show less speed modulation^32,34,35^, depend on cholinergic inputs^32,36-38^, and has been associated with emotions^39,40^, environmental novelty^30^, OLM interneuron activity in the ventral hippocampus^41^, and sensory processing^42^. In this respect, our cue-rich treadmill might have recruited lower theta frequencies because of the large engagement of visuo-tactile sensory systems. However, the necessity of path integration might explain the concurrent emergence of higher theta frequencies and the broader spectrum of grid cells’ theta frequencies in the path integration task.

Coinciding with this broadened frequency range was a reduction in grid size. Our model indicated that both effect were consistent with a theta-interference mechanism. A plausible scenario is that CAN and interference mechanisms coexist and normally operate in synchrony, but become desynchronized during path integration in a cue-rich environment, with interference producing shorter periodicities than the CAN. This mismatch could account for the altered grid coherence and short-timescale interactions observed in our task.

In summary, our results highlight the capacity of MEC networks to dissociate from landmarks and realign to rewards, enabling flexible support for goal-direct navigation. They further suggest that CAN and interference mechanisms of grid coding normally cooperate, but can diverge under the atypical requirement of path integration in cue-rich environments. Elucidating how these mechanisms interact across navigation regimes will be an important direction for future work.

## Materials and Methods

### Animals

All experiments were approved by the Institutional Animal Care and Use Committee of the Korea Institute of Science and Technology and conformed to the *Guide for the Care and Use of Laboratory Animals* (NRC, 2011). Male C57BL/6 mice (<6 months old) were used. Mice were house 2-5 per cage in a vivarium with 12h light/dark cycle.

### Preparation for head fixation

Surgical procedures followed^43^. Mice were anesthetized with isoflurane (3% induction, 0.5–1.5% maintenance, 1.5L/min airflow; Hana Pharm Co. Ltd., Korea). Two small screws were implanted above the cerebellum as reference and ground electrodes. The skull surface was covered with dental acrylic (Super-bond C&B kit, Sun Medical, Japan). A 3D-printed plastic head-plate^44^ (44Chung et al., 2017) was attached using dental acrylic, UV light-curing resin (Metafil Bulk Fill Low-Flow A2, Sun Medical, Japan) and self-curing resin (Vertex Self-Curing, Vertex Dental, Netherlands).

### Behavioral apparatus

The treadmill setup has been described previously^45,46^. It consisted of a 230 × 5 cm velvet belt carrying three pairs of visual-tactile cues. The belt was mounted on 3D-printed wheels and driven voluntarily by the mouse. Water rewards were delivered via a lick port with an LED and photosensor to detect licks. A microcontroller (Arduino Uno, Arduino SRL, Italy) monitored treadmill motion, controlled reward delivery and synchronized behavioral signals with electrophysiology.

The open field was a 90 × 90 cm white acrylic arena with 40-cm-high walls and three fixed visual cues. Illumination was <20 lux. Behavior was monitored using an infrared camera (acA130-60gmNIR, Basler AG, Germany) and EthoVision XT (Noldus, Netherlands).

### Behavioral protocol

Mice were handled for 10 min/day for 3 days before surgery. After a 7-day recovery period, they were water-restricted (1 ml/day) and trained to run on the treadmill for 30 min/day. Rewards were initially delivered after 40 cm, then increased to 65 and 105 cm once mice successfully completed >25 trials per session. Training continued at 105 cm was until lick rates in the 5-cm window before reward exceeded 2 SD above baseline. Trained mice performed 50-150 trials in 30 min. Water intake from the treadmill was supplemented to maintain ≥1ml of water/day. Mice were also trained to forage for scattered potato chip crumbs in the open field for 10 min/day.

### Silicon probe implantation

Mice were anesthetized with isoflurane and a craniotomy was made above the right MEC (3.2 mm lateral to midline, 0.5–0.8 mm anterior to the transverse sinus). A 64-channel silicon probe (Neuronexus, Buzsaki64sp), mounted on a custom microdrive^44^ and coated with DiI (Life Technologies) was lowered 0.8–1 mm with a 5° posterior tilt. The microdrive was secured to the skull with dental cement, the craniotomy sealed with bone wax/mineral oil, and protect with a 3D-printed cap^44^.

### Recording procedures

Broadband data were acquired at 30 kHz using an Intan RHD 2000 system and custom Labview software. Mice were recorded sequentially in the open field (25 min, free foraging) and treadmill (30 min, head-fixed). On the treadmill, mice performed either the PI task, the cue task, or both (cue task following 25 PI trials). The probe was advanced 25-100 µm daily to sample new units. Open-field firing maps were used to detect grid cells.

### Histology

At the end of experiments, mice were perfused with 4% paraformaldehyde. Brains were post-fixed overnight, sectioned sagittally (100 µm), and mounted with DAPI-containing Vectashield. For one mouse, parvalbumin immunolabeling was performed to delineate the postrhinal cortex^47-49^. Sagittal sections (50 µm) were incubated in PBST with 5% normal donkey serum (1 h, RT), then with chicken anti-parvalbumin antibody (1:2000, Synaptic Systems) overnight at 4 °C. After PBST washes, sections were incubated with Alexa Fluor 488 goat anti-Chicken IgY (Thermo Fisher) for 1 h, washed and mounted.

### Data acquisition and spike sorting

Open-field and treadmill data from the same day were concatenated. Signals were high-pass filtered (>300 Hz) for spikes and low-pass filtered (<500 Hz) and downsampled (1 kHz) for LFPs. Spikes were clustered with automatic algorithms (Mountainsort^50^) and refined manually in MATLAB using autocorrelograms, cross-correlations and cluster isolation statistics^51,52^.

### Firing rate maps

Spike trains and position data were binned at 10 ms. Firing rates were calculated in 1 cm^2^ (open field) or 1 cm (treadmill) bins by dividing spike counts by occupancy. Maps were smoothed with a Gaussian kernel (half-width = 10 cm).

### Grid score

Gridness was assessed in the open field using spatial autocorrelograms^21^. Grid scores were defined as the minimum correlation at 60°/120° rotations minus the maximum correlation as 30°/90°/150°. Cells with scores >0.3 were classified as grid cells.

To test for predictive coding^22^, we recomputed grid scores after shifting positions by offsets (-25 to +25 cm, 1-cm steps). The offset maximizing gridness was taken as predictive distance, and these maps were used in subsequent analyses.

### Border score

Border scores were computed following Solstad et al^2^. Fields were defined as contiguous pixels >30% peak rate covering >200 cm^2^. Coverage (c_M_) was the fraction of wall pixels contacted by the field with maximal overlap. Mean distance (d_m_) was the normalized, firing-rate-weighted distance to the nearest wall divided by 45 (half arena length). Border score = (c_M_ - d_m_) / (c_M_ + d_m_) and ranged from -1 (central fields) to +1 (perfect wall alignment). Cells were classified as border cells if scores >0.5, field stability >0.5, and preferred-wall scores were at least twice higher than others.

### Correlation matrices

For each trial, population vectors were constructed by concatenating simultaneously recorded firing maps. Pearson’s correlations between trial vectors yielded correlation matrices. Behavioral correlation matrices were constructed similarly using concatenated normalized speed and lick profiles. Trials were clustered with hierarchical clustering (MATLAB linkage/cluster functions; maximum clusters = N/5).

### Spatial coherence between grid cells

Coherence between grid maps was quantified as magnitude-squared coherence between vectorized maps. Power spectral densities of individual cells (*P*_*xx*_(*f*),*P*_*yy*_(*f*)) and cross-power spectral density *P*_*xy*_(*f*) were obtained via FFT, with *f* (cm^-1^) denoting the spatial frequency (Cycle = 1/*f*). The coherence was then 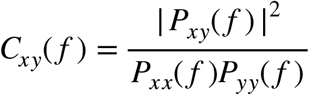

### Simulation of grid cells for linear trajectories through two-dimensional grid space

We simulated a module of 30 grid cells with a grid scale of 47.5 cm, a tilt angle of 0.18π, and randomly assigned phase offsets between 0 and 2π. Each cell’s two-dimensional activity map was computed by summing three sinusoidal planes rotated by 60° relative to one another, followed by thresholding and exponential filtering^23^. Linear activity patterns were obtained by intersecting 105-cm trajectory vectors with the 2D maps. In three sets of simulations, 10 trajectory vectors were generated per condition, incorporating either random rotation angles (rotation), x-axis offsets (collinear shifts), or y-axis offsets (parallel shifts).

### Grid size in the treadmill

Within each trial, firing rate vectors were normalized (0–1). Peaks were defined as local maxima >0.4, separated by ≥10 cm, with adjacent troughs <0.3. Distances between peaks were histogrammed (bin = 3 cm), and the peak of this distribution was taken as grid size.

### Theta frequency of local field potentials

LFPs were z-scored, detrended, and power spectra computed with FFT. Spectra were smoothed (300-point window), corrected for 1/f, and z-scored. The peak within 5–10 Hz was taken as theta frequency (median across channels).

### Theta frequency of grid cells

Spike autocorrelograms (±700 ms, 10-ms bins, speed > 5 cm/s) were fit in MATLAB with the equation *y*(*t*) = [*a* cos(*wt*) + *b*] · exp(−*t* /*τ*) + *c* where t is the autocorrelogram time variable and *a, w, b, c*, and *τ* are the fit parameters^53^. Cell theta frequency f = *w*/2π.

### Interference model

We implemented an oscillatory interference model with competitive inhibition. Grid cells (nGC = 100) received inputs from two groups of oscillators (n1 = 10, n2 = 10) with distinct frequencies (f1, f2) and random phase delays (0.01–0.1). Inputs were weighted randomly (gamma distribution, shape = 0.005). Simulations ran for 100 s at 1-ms resolution. Grid cell excitation was the weighted input sum, inhibition was proportional to network activity, and firing was E–I if >0, else 0^46^.

Theta frequency was derived from inter-peak intervals, and grid size from autocorrelograms implemented on firing rate envelops. Frequencies f1 and f2 were optimized via Nelder–Mead (MATLAB *fminsearch*) to match empirical mean (µ) and variance (σ) of theta distributions. Target values: PI µ=8.3 Hz, σ=0.79 Hz; Cue µ=7.9 Hz, σ=0.55 Hz; OF µ=9.1 Hz, σ=0.47 Hz. Solutions: PI f1=7.5 Hz, f2=9.4 Hz; Cue f1=7.4 Hz, f2=8.7 Hz; OF f1=8.4 Hz, f2=9.5 Hz.

### Short time scale interactions between grid cells

Spike cross-correlograms (±700 ms,10-ms bins, speed >5 cm/s) were computed. The short-time scale interaction (STSI) index = (*a* − *b*)/ max(*a, b*), where a is mean spike count in -10:10 ms window, and b is mean spike count in -400:-50 and 50:400 ms windows.

## Author contribution

SK and SR designed the project. SK performed experiments and analyses. SK and SR performed modeling and wrote the manuscript.

## Acknowledgments

This work was supported by National Research Foundation of Korea (NRF Grant # 2021R1A2C30055) and the Korea Institute of Science and Technology Institutional Program (Project # 2E33701) to SR, and Japan Society for the Promotion of Science (KAKENHI Grant # 24K18243) to SK.

