## Supplementary Figures for "Grid cells encode reward distance during path integration in cue-rich environments"

### Supplementary Materials for:

Grid cells encode reward distance during path integration in cue-rich environments

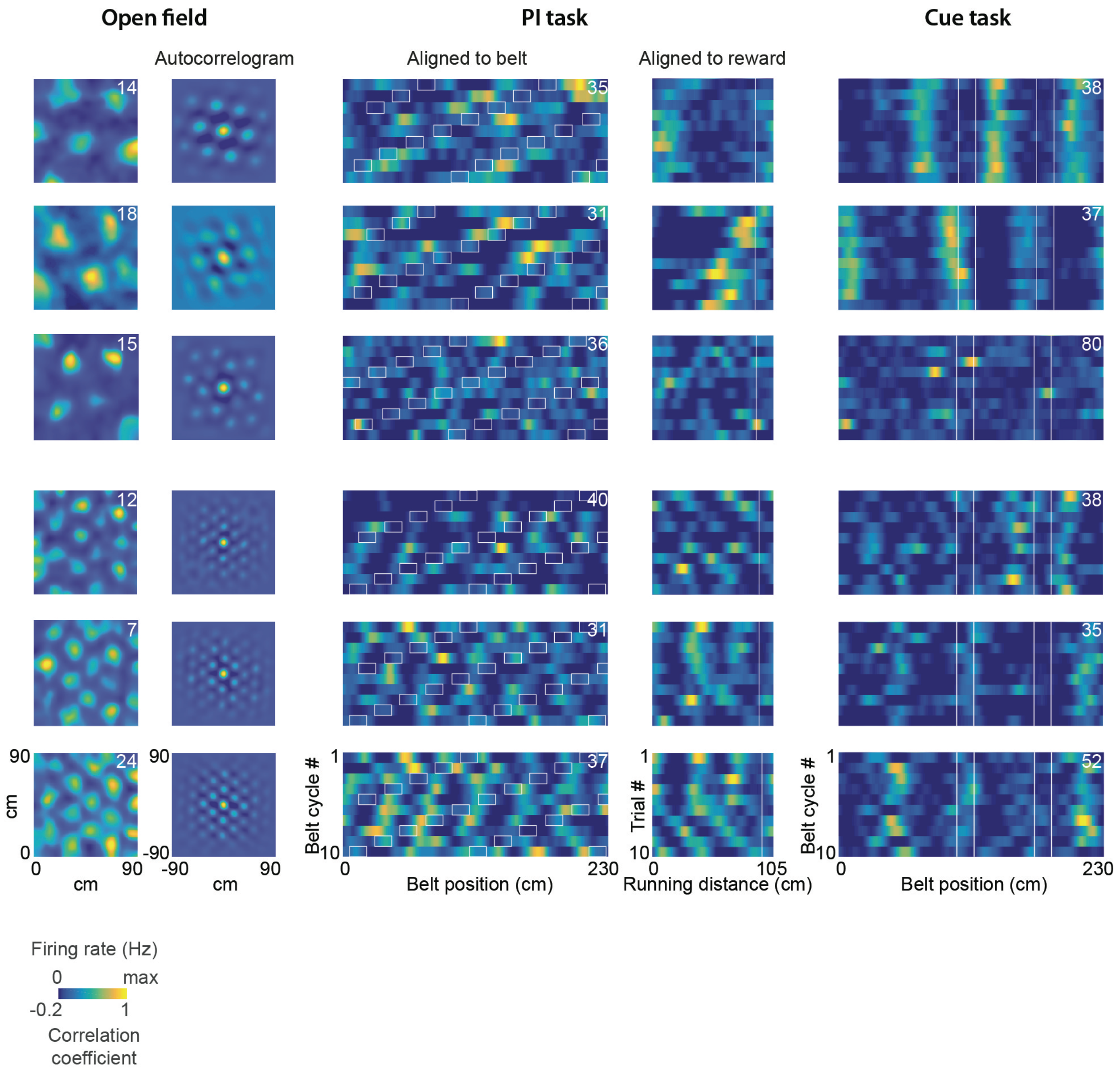


**Supplementary Figure 1**. Example grid cells recorded consecutively in open field, PI, and cue tasks. Rows show individual cells; displays as in Fig. 2.

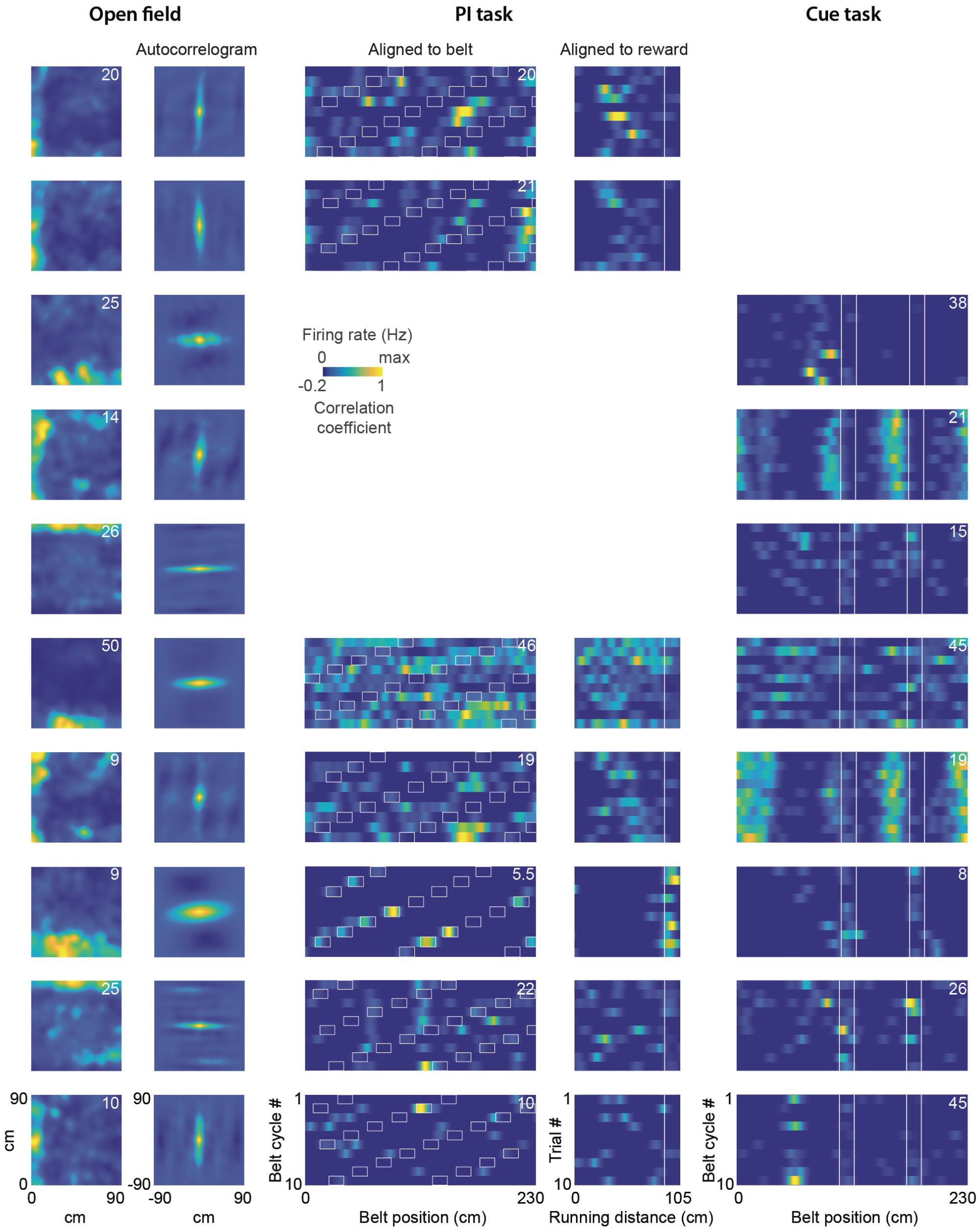


**Supplementary Figure 2**. Example border cells recorded across tasks, displayed as in Fig. 2. Bottom five cells were recorded consecutively in open field, PI, and cue tasks.

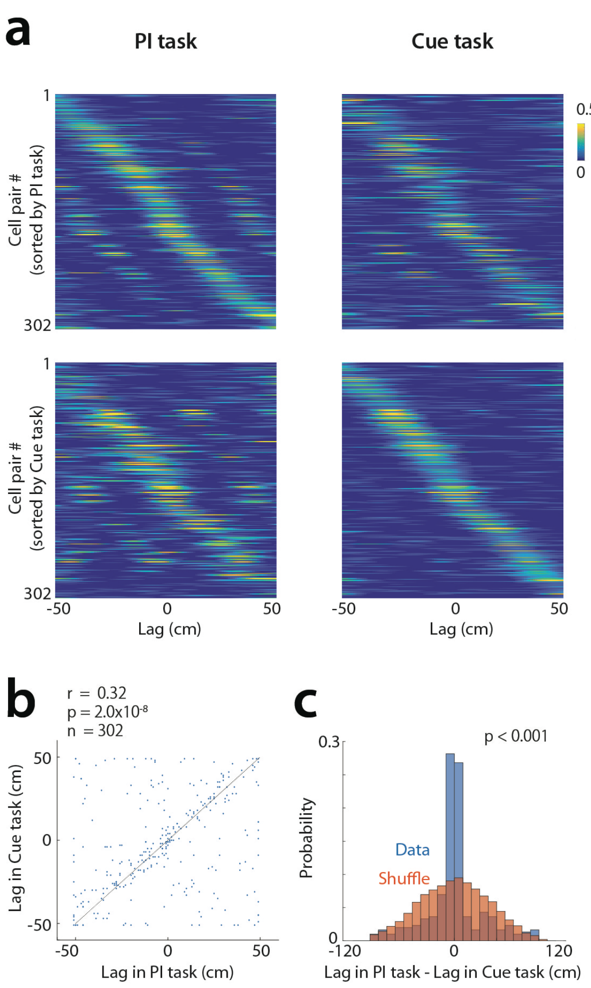


**Supplementary Figure 3**. Preservation of grid cell relationships across PI and cue tasks.

a, Spatial cross-correlograms (CCGs) for grid cell pairs recorded in PI (left) and cue (right). Pairs sorted by peak position in PI (top row) or cue (bottom row). Diagonal bands indicate preserved spatial relationships.

b, Cell–cell spatial shifts in cue versus PI tasks (n = 302 pairs; Pearson correlation).

c, Change in cell–cell spatial shifts across tasks (blue) compared to shuffled distributions (orange, 1,000 shuffles; n = 302 pairs; Wilcoxon signed-rank test on absolute values).

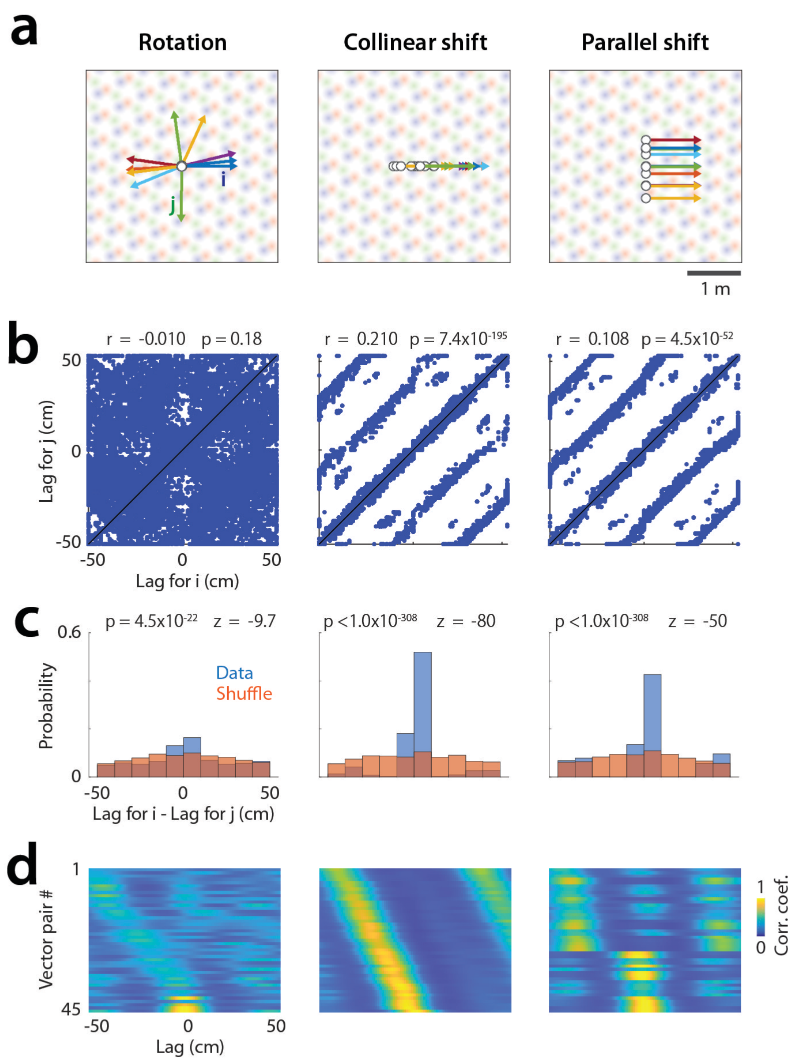


**Supplementary Figure 4**. Collinear trajectory shifts account for grid cell variability in the PI task.

a, Three trajectory vector alterations tested: rotation (left), collinear shift (middle), and parallel shift (right). Arrows, vectors (105 cm) used to generate firing rates for 30 model grid cells across 10 trajectories.

b, Spatial shifts between cell pairs across trajectory vectors, by alteration type (n = 19,575 trajectory pairs; Pearson correlation). Rotation does not reproduce Fig. 4e patterns.

c, Change in cell–cell spatial shifts across trajectories (blue) versus shuffled identities (orange, one shuffle per trajectory; n = 19,575 pairs; Wilcoxon rank-sum test).

d, Spatial cross-correlograms between population vectors of trajectory pairs. Parallel shifts fail to reproduce Fig. 4c–d patterns.

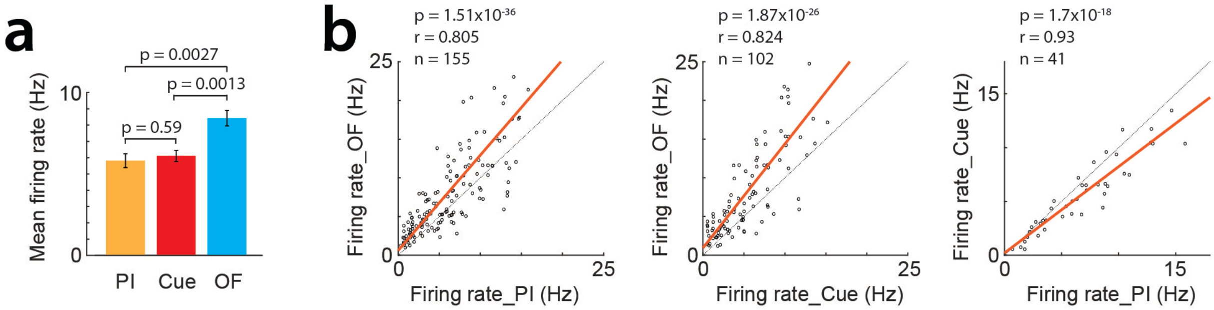


**Supplementary Figure 5**. Gird cell firing rates across tasks.

a, Firing rates (mean ± s.e.m., n = 155 PI, 102 Cue, 216 OF cells; Wilcoxon rank-sum test).

b, Firing rates in pairwise task comparisons (n = 155 PI vs OF, 102 Cue vs OF, 41 PI vs Cue cells; Pearson correlation).

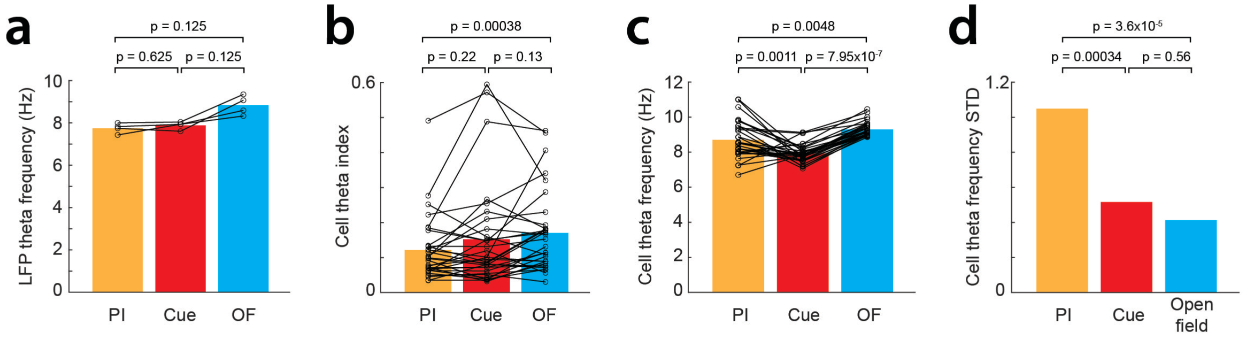


**Supplementary Figure 6**. Theta index and frequency in consecutively recorded sessions.

a, LFP theta frequency (mean ± s.e.m., n = 3 sessions; Wilcoxon signed-rank test).

b, Grid cell theta index (mean ± s.e.m., n = 32 cells; Wilcoxon signed-rank test).

c, Grid cell theta frequency (same cells as in b).

d, Standard deviation of grid cell theta frequency (same cells; Levene's test).
